# Euclidean distance as a measure to distinguish ventral and dorsal white matter connectivity in the human brain

**DOI:** 10.1101/053959

**Authors:** Philipp Kellmeyer, Magnus-Sebastian Vry

## Abstract

Fiber tractography based on diffusion tensor imaging (DTI) has become an important research tool for investigating the anatomical connectivity between brain regions in vivo. Combining DTI with functional magnetic resonance imaging (fMRI) allows for the mapping of structural and functional architecture of large-scale networks for cognitive processing. This line of research has shown that ventral and dorsal fiber pathways subserve different aspects of bottom-up- and top-down processing in the human brain.

Here, we investigate the feasibility and applicability of Euclidean distance as a simple geometric measure to differentiate ventral and dorsal long-range white matter fiber pathways tween parietal and inferior frontal cortical regions, employing a body of studies that used probabilistic tractography.

We show that ventral pathways between parietal and inferior frontal cortex have on average a significantly longer Euclidean distance in 3D-coordinate space than dorsal pathways. We argue that Euclidean distance could provide a simple measure and potentially a boundary value to assess patterns of connectivity in fMRI studies. This would allow for a much broader assessment of general patterns of ventral and dorsal large-scale fiber connectivity for different cognitive operations in the large body of existing fMRI studies lacking additional DTI data.

## 1. Introduction

In recent years, fiber tracking based on diffusion tensor imaging (DTI) and functional magnetic resonance imaging (fMRI) have allowed for investigation of large-scale white matter pathways for cognitive processing in different domains (Vry et al., 2012; Saur et al., 2008; Umarova et al., 2010). This research has demonstrated that ventral and dorsal fiber systems subserve different aspects of cognitive processing in a variety of domains like vision, attention, language and the motor system. In the context of language processing, this line of research has demonstrated that, in addition to the classic dorsal pathway (via the fiber system of the arcuate fascicle and superior longitudinal fascicle), a ventral pathway via the fibers of the middle longitudinal fascicle and the extreme capsule connects parietal and posterior temporal areas for speech perception, comprehension and phonological processing with inferior frontal areas for articulation (Kellmeyer et al., 2013; Saur et al., 2010, 2008; Frey, Campbell, Pike, & Petrides, 2008). Overall, this research has produced considerable insight into the structural white matter architecture, specific patterns of large-scale connectivity and the neuronal implementation of different cognitive systems in the brain.

In-vivo mapping of human fiber pathways depends on DTI measurements, which have only become widely available for cognitive neuroscience research about a decade ago. However, we have a large body of functional imaging studies from before the advent of DTI, in which the patterns of structural connectivity between the brain regions identified with fMRI (or PET) remain unexplored. Thus, it would be useful if the current knowledge of the human fiber system from DTI-based fiber tracking could be harvested to retrospectively infer patterns of connectivity between brain coordinates in the absence of DTI data. This would allow for much larger meta-analyses on patterns of large-scale connectivity across a variety of cognitive domains, and therefore would increase the aggregated evidence on large-scale networks for cognitive processing based on functional neuroimaging.

Here, we explore systematic differences in Euclidean distance (ED) in studies that have used DTI-based probabilistic fiber tracking to test it as a simple geometric measure to distinguish between ventral and dorsal pathways for different aspects of cognitive processing. Our aim is to investigate whether the anatomical segregation into ventral and dorsal fiber pathways, demonstrated in recent studies by probabilistic fiber tracking, can be captured by a simple measure of spatial distance. Differentiating ventral and dorsal connectivity between brain coordinates by Euclidean distance would provide a simple tool for assessing patterns of connectivity in the brain even in the absence of DTI data, as in a large body of fMRI studies.

Euclidean distance has been previously shown to relate to functional connectivity between brain regions in resting state fMRI in the context of psychiatric disorders. In a study in patients with childhood-onset schizophrenia, the differences between subjects in the topology of functional networks in the resting-state was reflected by variation in ED between connected regions (Alexander-Bloch et al., 2013). Another study which compared resting-state fMRI between elderly patients with depression and controls showed increased ED between connected brain regions in the depressed group (Bohr et al., 2013). With respect to long-range anatomical connectivity via white matter fiber pathways in the brain, however, no geometrical relationships, power laws or allometric principles based on in-vivo DTI imaging data is available so far to our knowledge.

Here we analyzed Euclidean distances between coordinates in parietal and inferior frontal brain regions for which the pattern of ventral or dorsal connectivity has been demonstrated previously by DTI-based probabilistic fiber tracking. We chose probabilistic fiber tracking because we have substantial experience in developing and applying this fiber tracking method (Kreher et al., 2008). In this previous work, probabilistic fiber-tracking investigated ventral and dorsal fronto-parietal fiber pathways in the context of language processing (Kellmeyer et al., 2013; Saur et al., 2008, 2010), motor imagery (Hamzei et al., 2016; Vry et al., 2012), attention (Suchan et al., 2014; Umarova et al., 2010), and mental arithmetic (Klein, Moeller, Glauche, Weiller, & Willmes, 2013) we observed that ventral fronto-parietal fiber pathways generally seem to travel much longer distances in the brain than dorsal fronto-parietal pathways. However, experiments on perception show that visual estimation of geometric distance between two points often results in an overestimation of the distance, with relative errors increasing with the length of curvilinear paths (that were used as visual cues, called the “detour effect”) (Faineteau, Gentaz, & Viviani, 2003). Based on our observation from these tractography studies, we hypothesize that the ventral pathway connecting parietal and inferior frontal cortex is on average longer than the dorsal pathway. To test this hypothesis, we analyze systematic geometric differences in ventral and dorsal connectivity between parietal and inferior frontal regions quantitatively by measuring Euclidean distance in 3D coordinate space in these studies.

## 2. Materials and methods

To test this approach, we confined the analysis to two well characterized regions in terms of fiber connectivity, the parietal lobe and the inferior frontal gyrus (opercular and triangular parts). As we have studied white matter fiber connectivity using probabilistic fiber-tracking in a variety of cognitive domains and have a good understanding of the methodology of these measurements, we confined this investigation to these studies using probabilistic fiber tracking (see **Table 1**).

### 2.1. Identification of coordinates

For the list of studies that were used for identifying the coordinates and their respective connectivity pattern from the fiber tracking see **Table 1**. All studies that were included in the analysis reported brain coordinates in the Montreal Neurological Institute (MNI) coordinate space.

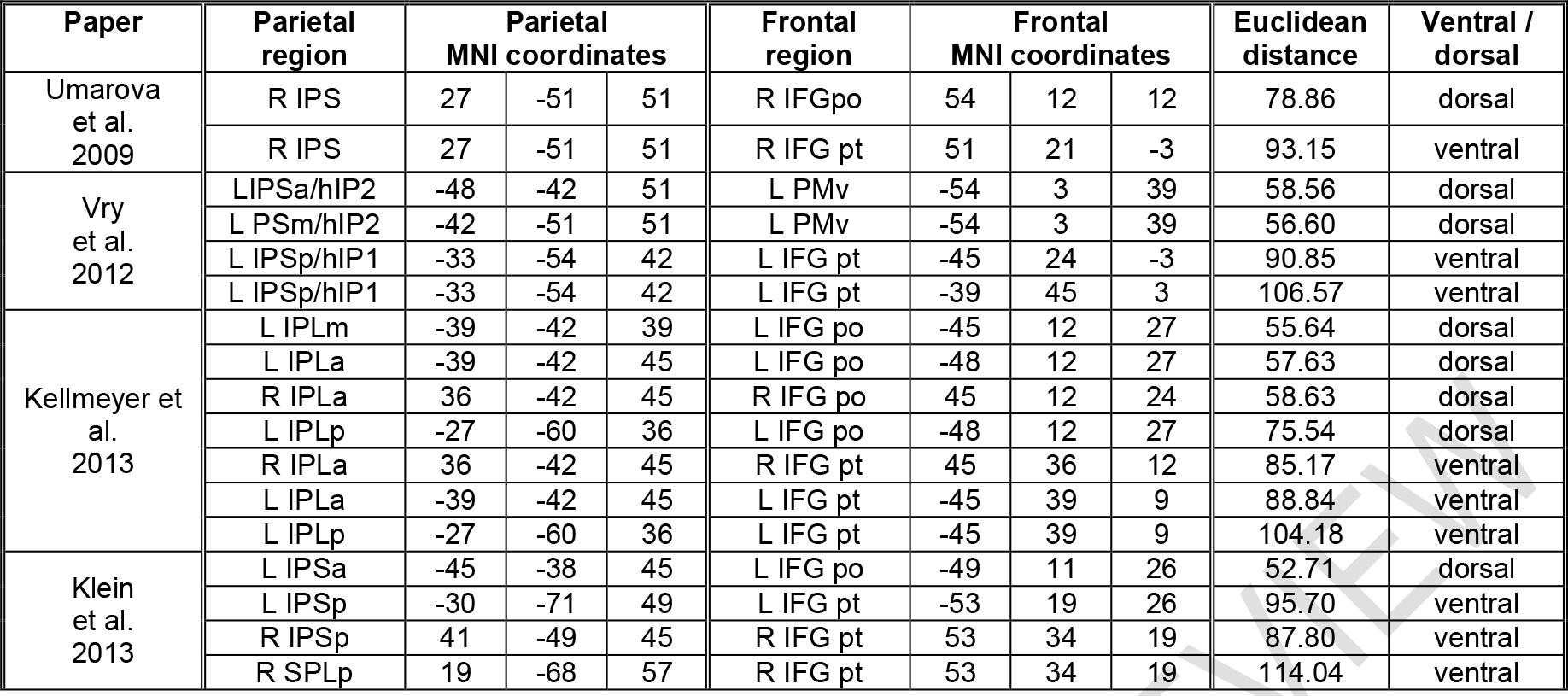

### 2.2. Measuring geometrical distance in 3D coordinate space

We used Euclidean distance (ED) 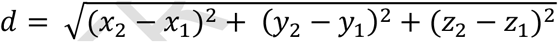 to measure the distance between two coordinates in three-dimensional MNI space, located in the inferior parietal and inferior frontal cortex respectively, for which connectivity has been demonstrated previously by DTI-based probabilistic fiber tracking. Fiber pathways from the tracking results were either classified as dorsal or ventral in accordance with the classification from the respective studies (Fig. 1).

**Fig. 1.**
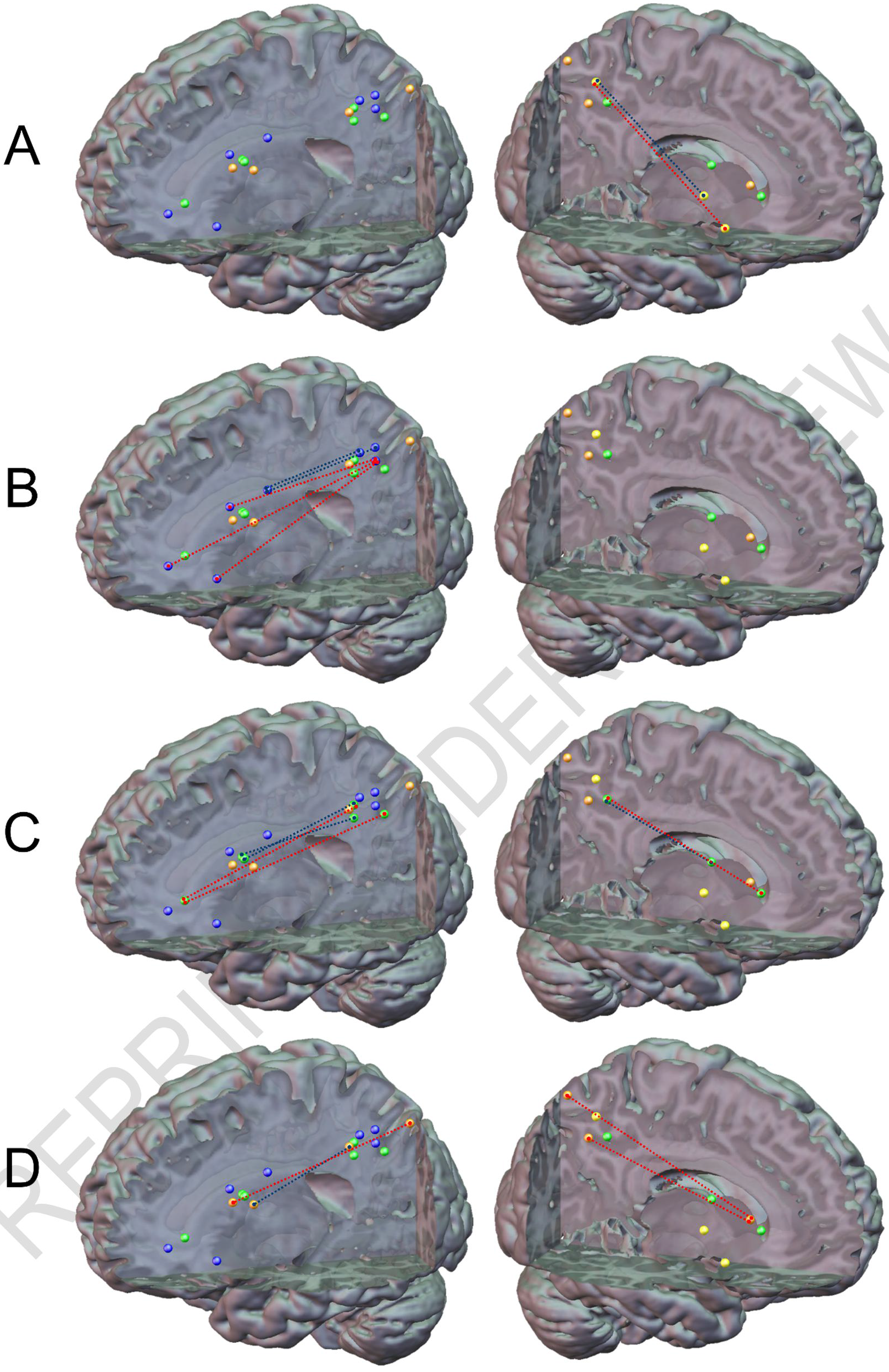
Patterns of dorsal and ventral connectivity from the studies used for measuring Euclidean distance: Panels (A)-(D) show the different Studies with each panel including the seeds from all studies. (A) Umarova et al. 2009 (yellow spheres), (B) Vry et al. 2012 (blue spheres), (C) Kellmeyer et al. 2013 (green spheres), (D) Klein et al. 2013 (orange spheres). Blue dotted line = dorsal pathway; red dotted line = ventral pathway. For MNI coordinates (see **Table 1**)

### 2.3. Comparing mean Euclidean distance between dorsal and ventral connections

The null-hypothesis, that Euclidean distance between dorsal and ventral pathways connecting parietal and inferior frontal cortex does not significantly differ, was addressed in a two-sample t-test comparing the mean ED between pathways classified as dorsal or ventral by DTI-based probabilistic fiber tracking.

## 3. Results

### 3.1. Euclidean distance between ventral and dorsal fronto-parietal pathways

The two-sample t-test comparing median Euclidean distance (ED) between brain coordinates in parietal and inferior frontal cortex showed that ED for coordinates connected by ventral pathways (*N*=9, mean 96.26, standard deviation 9.84, standard error 3.28) are significantly (p<0.0001) longer than for dorsal pathways (*N*=8, mean 61.77, standard deviation 9.75, standard error 2.43) (Figure 2).

**Fig. 2.**
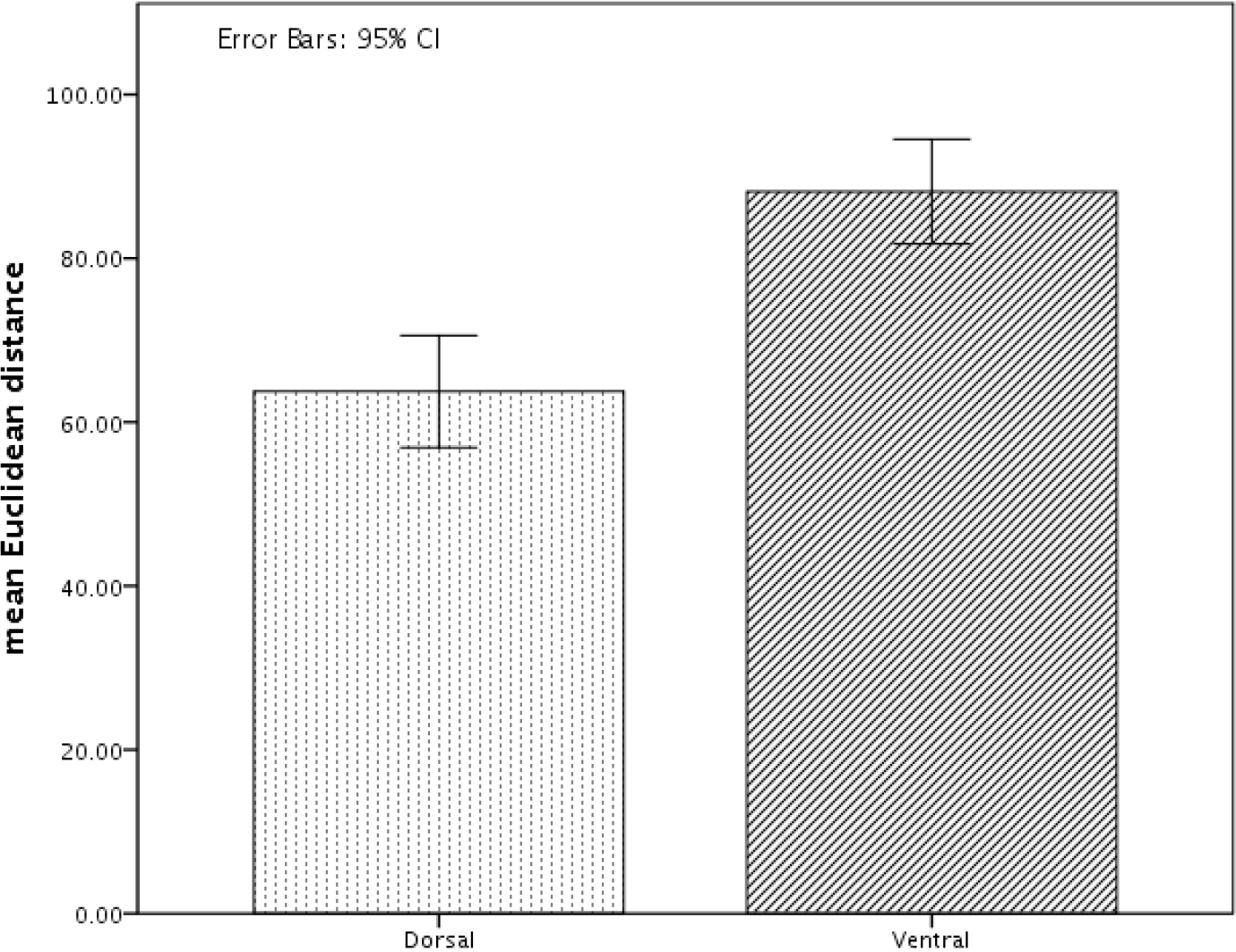
Mean Euclidean distance for dorsal and ventral pathways between the parietal and inferior frontal coordinates from **Table 1**

## 4. Discussion

In this sample of DTI-based probabilistic fiber tracking studies that assessed the large-scale pathway architecture of brain networks engaged in different cognitive domains, we found a significant difference for Euclidean distance to distinguish ventral from dorsal pathways between frontal and parietal regions.

Therefore, we propose Euclidean distance as a feasible albeit simple measure to distinguish ventral from dorsal connectivity between these brain regions, applicable in the absence of DTI data.

One explanation for this pattern could be that ventral parieto-frontal connections tend to connect to more anterior-inferior subregions in IFG (pars triangularis, pars orbitalis), whereas dorsal pathways tend to connect to more posterior-superior subregions (pars opercularis). This raises the question, whether there is an anatomical partition within IFG that determines ventral or dorsal connectivity to posterior brain regions like parietal and temporal cortex.

To replicate and validate the findings here and to address this question, we propose the following steps for further analyses: First, a voxel-by-voxel connectivity analysis between defined volumes in parietal cortex (comprising, for example, inferior and superior parietal lobule) and inferior frontal cortex (comprising inferior frontal gyrus, pars opercularis triangularis and orbitalis) could be performed to map the complete ventral/dorsal connectome between these two volumes.

Previous research with such a DTI-related connectivity-based parcellation (Johansen-Berg & Rushworth, 2009) approach has already shown the pattern of connectivity for inferior frontal cortex (Anwander, Tittgemeyer, Cramon, Friederici, & Knösche, 2007), orbitofrontal cortex (using fMRI) (Kahnt, Chang, Park, Heinzle, & Haynes, 2012), and inferior parietal cortex (Ruschel et al., 2014) in separate studies. However, no voxel-to-voxel whole connectome between parieto-frontal regions is available thus far. If such a whole region-to-region connectome was available, the ED between all coordinates for which either ventral and/or dorsal connectivity was found by fiber tracking could be measured. Then the mean ED could be compared between ventral and dorsal pathways and the boundary ED value that reliably (for example, >95% of cases) distinguishes between ventral and dorsal connectivity could be computed. In addition, this data could then be used to delineate a possible anatomical boundary at which fronto-parietal connectivity segregates between ventral and dorsal pathways.

This measure would be a particularly useful tool for large-scale meta-analysis of fMRI studies which did not measure additional DTI data. In such a meta-analysis, one could address the question whether the network involved in particular cognitive operations, e.g. speech repetition based on semantic versus phonological features, can be assigned with high probability to different pathways according to the MNI coordinates that were identified in fMRI studies on the topic. Information on the structural connectivity of “active” brain regions in neuroimaging studies is of interest for both cognitive and clinical neuroscientists.

For cognitive functions that operate via long-range dorsal and ventral fiber tracts, computational efficiency may be influenced by a number of anatomical factors and constraints. It has been shown, for example, that the average conduction delay in axonal transmission scales with the average length of long-range connections and average axon diameter (Herculano-Houzel, Mota, Wong, & Kaas, 2010). In the healthy brain, moreover, differences in domain-independent cognitive processing speed correspond to the length of white matter pathways connecting regions involved in the tasks used (Behrman-Lay et al., 2014).

Synthesizing findings from functional and structural imaging studies, dorsal and ventral pathways seem to indeed subserve different functions. In the case of language, for example, the ventral temporo- and parieto-frontal pathways is more involved in “higher-order” aspects of language—like syntactic or lexical-semantic processing (Saur et al., 2008). The same seems to be the case for cognitive aspects of the motor system like imitation of gestures and pantomime (Hamzei et al., 2016). More generally, it has been proposed that the ventral pathway might be used for processing that is time-independent and related to semantic aspects whereas the dorsal pathway is involved in sequential processing in a time-sensitive manner for rapid sensorimotor integration (Kellmeyer et al., 2013; Rijntjes, Weiller, Bormann, & Musso, 2012).

From a clinical perspective, whether a particular pathway is significantly longer than another pathway and/or may traverse regions that are more prone to damage (e.g. due to differences in territorial vascular supply) may also entail differences in neurological vulnerability. These differences in vulnerability between ventral and dorsal pathways, in turn, may partly explain different clinical phenotypes of cognitive deficits after brain injury, for example the different aphasia syndromes (McKinnon et al., 2017; Kümmerer et al., 2013).

More generally, it might be interesting to investigate how geometric measures of connectivity, whether Euclidean distance or other measures, are useful metrics to differentiate patterns of fiber connectivity in other mammalian species, for example non-human primates for which neuroanatomical data in 3D coordinate space is available (Markov et al., 2013). This may give further insight into cross-species principles of white matter formation in nervous systems, particularly links between global principles like minimal wiring and more local organizing principles of neuronal circuitry and topographical maps (Rothschild & Mizrahi, 2015; Goñi et al., 2014; Perin, Berger, & Markram, 2011). Such information would also be important to link the global geometry of the white matter fiber system to the hierarchical organization of multimodal cortical areas (Zhang & Sejnowski, 2000).

Importantly, the precise relationship between ED and actual fiber length cannot be inferred from the analysis presented here. It may well be the case, for example, that the given pathways are optimal in terms of their distance under the given physical and anatomical constraints on the actual brain’s global shape, local curvature, folding and gyration, Furthermore, as the pathways are not necessarily monosynaptic connections, the ventral pathway is relayed in insular cortex for instance, we do not know how ED relates to whether a particular pathway is mono- or polysynaptic. In this respect, it could be of interest to investigate whether the relationship between Euclidean distance and patterns of ventral and dorsal connectivity may reflect some kind of underlying allometric scaling law for governing principles of long-range white matter fiber connectivity in the human—and more generally the mammalian—brain (Herculano-Houzel et al., 2010; Bush & Allman, 2003; Harrison, Hof, & Wang, 2002; Changizi, 2001; Zhang & Sejnowski, 2000; Gould, 1975). In connectomics research, a major challenge is to find commensurate solutions that satisfy both the realities of neurophysiological signaling time scales, neuroanatomical constraints and the desired (in terms of computational modeling) optimality of network communication under the given constraints of small- and large-scale wiring in the human brain (Seguin, Van Den Heuvel, & Zalesky, 2018; Sporns, 2011). Such a parsimonious measure of network geometry as ED could also be an important baseline against which to compare other methods of comparing the topographical organization of brain networks (Martínez & Chavez, 2018).

To this end, other modeling approaches based on non-euclidean, curvilinear measures of geometric distance between given coordinates may also provide important insight in on this question (Styner, Coradi, & Gerig, 1999). In terms of the underlying geometrical principles that may best describe long-range fiber connectivity, algorithms for DTI analysis based on Riemannian and geodesic distance measures have been used that could have advantages over the currently used voxel-based analyses (Yamin et al., 2019; Wang, Yap, Wu, & Shen, 2014; Hao, Whitaker, & Fletcher, 2011; Astola, Florack, & ter Haar Romeny, 2007).

Further studies should also address whether differences in Euclidean distance related to ventral and dorsal connectivity can also be found in connections between other brain regions (for instance fronto-temporal, occipito-temporal etc.). Moreover, DTI-based connectivity studies that use different tracking methods such as deterministic fiber tracking should also be examined in order to facilitate large-scale meta-analyses on the structure and function of dorso-ventral pathway systems based on fMRI studies.

## 5. Conclusions

Euclidean distance between brain coordinates could be a useful method to distinguish patterns of long-range white matter fiber connections in the brain, such as ventral and dorsal pathways. This measure might be particularly useful to infer patterns of ventral/dorsal connectivity from brain coordinates in neuroimaging studies in the absence of DTI data. This, in turn, could facilitate neuroimaging meta-analyses that include data from the many neuroimaging studies without concomitant DTI data from the subjects.

## Acknowledgement

We thank Prof Dorothee Saur (University of Leipzig, Medical Center) for valuable discussions on a previous draft of the paper.

## Research involving Human Participants and/or Animals

The research reported in this paper itself involved no experiments on human subjects and/or animals but consists of an analysis of previously published research papers. Please refer to the papers with respect to the research on human participants performed by the respective researchers.

## Ethical approval

The research reported in this paper itself involved no experiments on human subjects and/or animals, therefore no ethics approval was sought. The research papers analyzed here all reported ethical approval for the research with human participants performed by the respective researchers.

## Informed consent

The research reported in this paper itself involved no experiments on human subjects and/or animals, therefore procuring informed consent was not applicable here. The research papers analyzed here all reported to having obtained informed consent for the research with human participants performed by the respective researchers.

